# Lisen&Curate: A platform to facilitate knowledge tools for curation of regulation of transcription initiation in bacteria

**DOI:** 10.1101/2020.04.28.065243

**Authors:** Carlos-Francisco Méndez-Cruz, Martín Díaz-Rodríguez, Francisco Guadarrama-García, Oscar Lithgow-Serrano, Socorro Gama-Castro, Hilda Solano-Lira, Fabio Rinaldi, Julio Collado-Vides

## Abstract

The amount of published papers in biomedical research makes it rather impossible for a researcher to keep up to date. This is where machine processing of scientific publications could contribute to facilitate the access to knowledge. How to make use of text mining capabilities and still preserve the high quality of manual curation, is the challenge we focused on. Here we present the Lisen&Curate system designed to enable current and future NLP capabilities within a curation environment interface used in curation of literature on the regulation of transcription initiation in bacteria. The current version extracts regulatory interactions with the corresponding sentences for curators to confirm or reject accelerating their curation. It also uses an embedded metrics of sentence similarity offering the curator an alternative mechanism of navigating through semantically similar sentences within a given paper as well as across papers of a pre-defined corpus of publications pertinent to the task. We show results of the use of the system to curate literature in *E. coli* as well as literature in *Salmonella*. A major advantage of the system is to save as part of the curation work, the precise link for every curated piece of knowledge with the corresponding specific sentence(s) in the curated publication supporting it. We discuss future directions of this type of curation infrastructure.

## INTRODUCTION

Manual curated databases are a major resource to facilitate access to the ever-increasing biomedical knowledge described in scientific publications. Among the different efforts to gather knowledge of specific biological domains (See the special issue of databases in *Nucleic Acids Res*), we initiated close to 30 years ago the gathering of knowledge around regulation of transcription initiation in bacteria, in a review with 117 regulated promoters (Collado-Vides et al., 1991). This was the seed of what became later on RegulonDB (Huerta et al., 1998) which we have maintained since then, feeding also this knowledge into EcoCyc (REF).

The more than 1000 citations of RegulonDB through the years, indicate how useful this organized knowledge has been to research projects worldwide. Nonetheless, we are aware that there are several layers, so to speak of additional higher-level knowledge of the molecular biology and physiology of gene regulation in *E. coli* that are not captured by our curation work. Reading a paper and comparing that experience with the one of only looking at the objects and interactions supported by that paper present in RegulonDB, is a troubling exercise. The pieces of knowledge can be accurate, but a rich amount of material is left out. The argumentation of additional evidence, the connections of mechanisms and operon organization to the physiology of *E. coli*, the interpretation of the results, and additional questions to be solved, are all absent in RegulonDB or EcoCyc. A small part may be included in the notes added to objects, and in the rich summaries of TFs. Additional rich knowledge is present in reviews that experts publish about a variety of components of transcriptional regulation and its physiological interpretation. Of course, these reviews are usually not curated since they lack any specific new molecular or sequence element that our model behind the database contains and that we curate.

These observations point to a large number of exciting challenges that we have not even yet enumerated. Nonetheless, they motivated us to we call L-Regulon (L for “linguistic”) (See:https://www.biorxiv.org/content/10.1101/2020.04.26.062745v1), and the work here presented is the necessary curation infrastructure companion associated to it. Lisen&Curate is the name of this linguistic semantic enrichment curation infrastructure. One of the goals of Lisen&Curate is to offer an infrastructure that will save from the curation work the connections between curated objects and their precise sentence in the paper that supports its evidence. We also envision Lisen&Curate to offer curators natural language processing (NLP) capabilities that we have and will continue to develop to enrich the curation experience.

We developed Lisen&Curate to work with multiple bacteria. At present, the K 12 strain of *Escherichia coli* is available, and the strains LT2 (GCF_000006945.2_ASM694v2), SL1344 (GCF_000210855.2_ASM21085v2), ST4/74 (GCF_000188735.1_ASM18873v1000005845.2) and 14028s (GCF_000022165.1_ASM2216v1) of *Salmonella enterica subsp. enterica serovar Typhimurium*. Data of biological objects is retrieved from BioCyc PGDBs, and can be updated as Lisen&Curate stores them in a local database.

## LISEN&CURATE DESCRIPTION

Lisen&Curate is a Web system to assist curation of transcriptional regulation of bacteria that includes text mining tools and user interfaces to capture transcriptional regulatory information. We aim to improve curation work by offering automatically extracted and enriched source information for curators to make better and faster decisions. This is achieved due to the following Lisen&Curate tools, which are described in further subsections:

- Inter- and intra-document navigation. Lisen&Curate includes a text mining tool to interlink semantically similar sentences from the same article and from an article collection. Once a curator selects a sentence, similar sentences can be accessed using a navigation interface. This drives a curator to confirm an evidence or expand information of a transcriptional regulatory fact.
- Regulatory sentence detection. Lisen&Curate automatically detects regulatory sentences using a predictive model trained by ourselves with machine learning techniques. Then, curators see sentences with regulatory information highlighted to access relevant information directly.
- Regulatory interaction extraction. Lisen&Curate incorporates an in-house system to automatically extract regulatory interactions (RIs) between transcription factors and genes. This system finds out the regulatory effect: regulation, activation or repression. Extracted RIs are confirmed or rejected by curators utilizing specialized interfaces. If the RI is confirmed, this is stored in the Lisen&Curate database.

When curators find out and confirm evidence to register or modify transcriptional regulatory information of a biological object, they can open an interface to capture it. Lisen&Curate holds interfaces for genes, transcription units, promoters, regulatory interactions, proteins, proteins compounds, proteins features and terminators

An innovative functionality of Lisen&Curate is that each feature of the objects can be associated to some textual sources, that is, to fragments of articles where the evidence of that feature is found. The set of textual sources constitute an important resource of evidence and may help as training data to develop automatic methods to extract those features.

Lisen&Curate provides a complete working environment in a main interface to read scientific articles. This interface comprises all tools for curators to work with extracted and enriched information, and to filter sentences by biological entities. Also, it includes all interfaces for capturing data, which can be accessed as needed.

Following subsections contain additional information of Lisen&Curate features and tools, as well as some technical descriptions.

### Curation processes

A *curation process* is a working concept for Lisen&Curate comprising an aim of curation, a set of curators (one of them as a reviewer) and an article collection (Fig. 1a). This concept let the RegulonDB team organize the curation work by establishing well-defined curation goals and set of articles previously selected. Then, the first step to work with Lisen&Curate is to select or create a curation process. This is accessible through the option *Curation process* in the Welcome interface (Fig. 1b). This option displays an interface with the list of all the curation processes in the system, as well as the button to create a new curation process (Fig. 1c). A curation process must include an article collection (collection) previously processed. This collection comprises the articles to be worked by curators.

**Figure 1.**
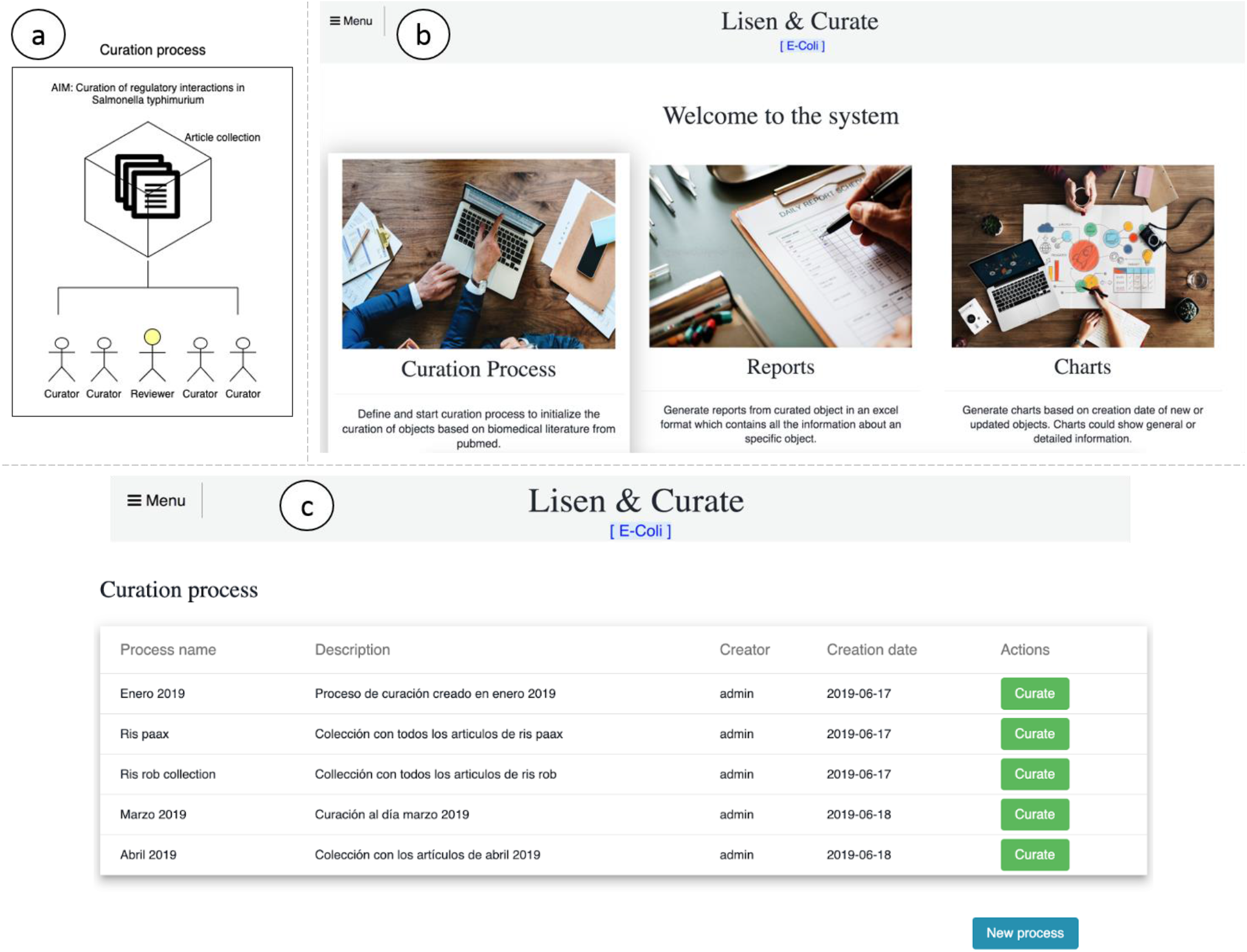
Curation processes in Lisen&Curate. (a) Graphical depiction of the concept of curation process. (b) Main interface of the system. (c) Curation process interface.

### Processing article collections

An *article collection* (*collection*) is a set of scientific articles associated with a name, previously processed to be displayed for Lisen&Curate. We implemented a pipeline to process article collections (Figure 2). In Step 1, we extracted text from PDF articles using a home-made tool. Output of this step are JSON documents with the content of the article and metadata. Our tool detects sections of the articles, which helps to visualize them in a structured way. In Step 2, extracted text undergoes some Natural language processing (NLP) tasks that covers sentence split, part of speech tagging, and dictionary-based named entity recognition (NER). Now, our dictionaries compris biological objects of *E. coli* and *Salmonella*. The output of this step is a JSON document per article with all this grammatical information.

**Figure 2.**
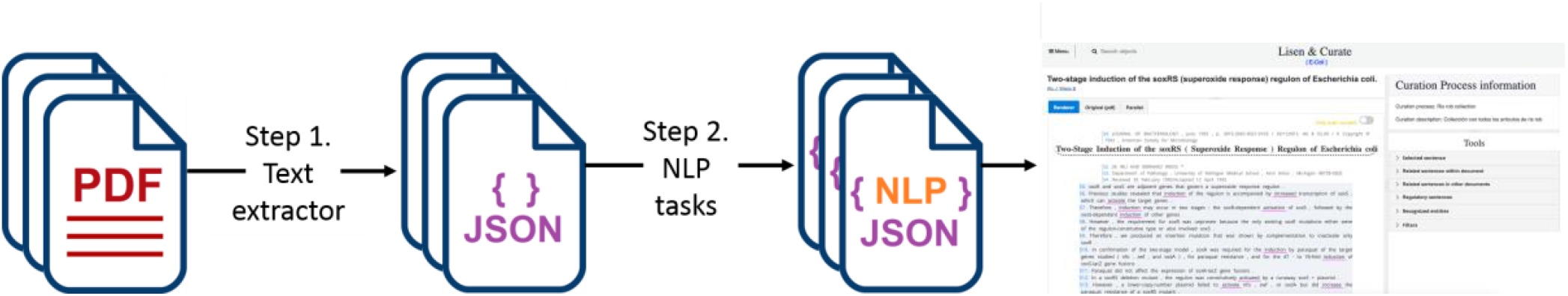
Processing article collections.

To create the collection in Lisen&Curate a two-step process is done:

1. Locating the files generated by the processing step into a directory.
2. Running a script for integrating the articles in Lisen&Curate and defining a collection name.

### Creating curation processes

Using graphical interfaces, users can create a curation process by choosing an existing collection (Fig. 3a). All articles that belong to the collection will be automatically included to the curation process, but users have the option to exclude some articles disabling the *Selected* button (Fig. 3b). Finally, curators and the reviewer are selected to work with the curation process, and the name and description of the curation process are also captured (Fig. 3c).

**Figure 3.**
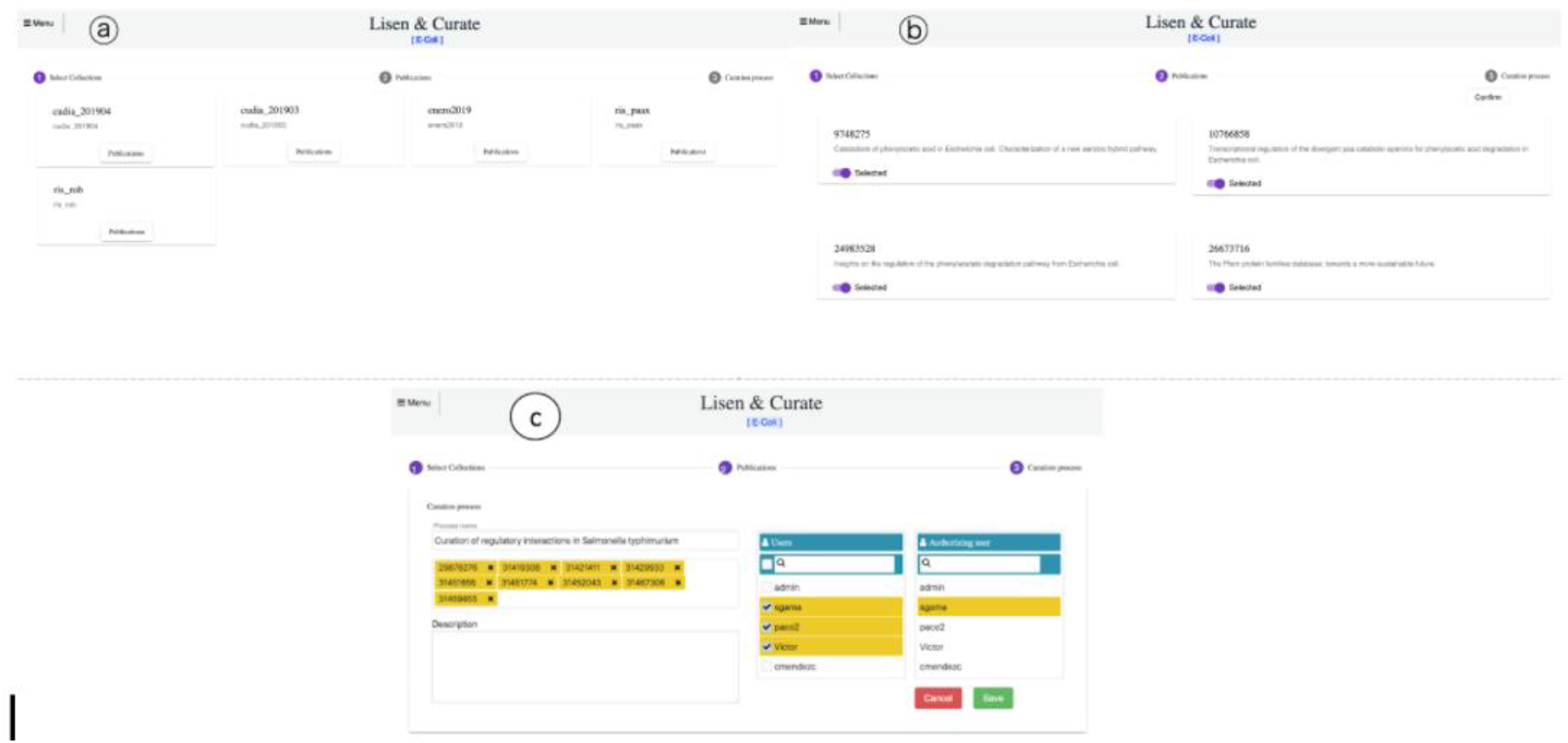
Creation of curation processes. (a) Selection of an available collection. (b) Confirmation or elimination of individual articles from selected collection. (c) Capturing name, description, curators and reviewer for a curation process.

### Working environment

#### Main interface

The main interface of the working environment is split into following resizable sections:

- Title: Display the title and the authors of the article (Fig. 4a).
- Article content: Displays the article as text, pdf or both formats (Fig. 4b).
- Meta information: Provide extra information about the article as well as the curation process related (Fig. 4c).
- Tools: Set of toolboxes to assist curation work (Fig. 4d).
- Interfaces for capturing data: Interfaces to capture and store regulatory data associated to biological objects (Fig. 4e).

**Figure 4.**
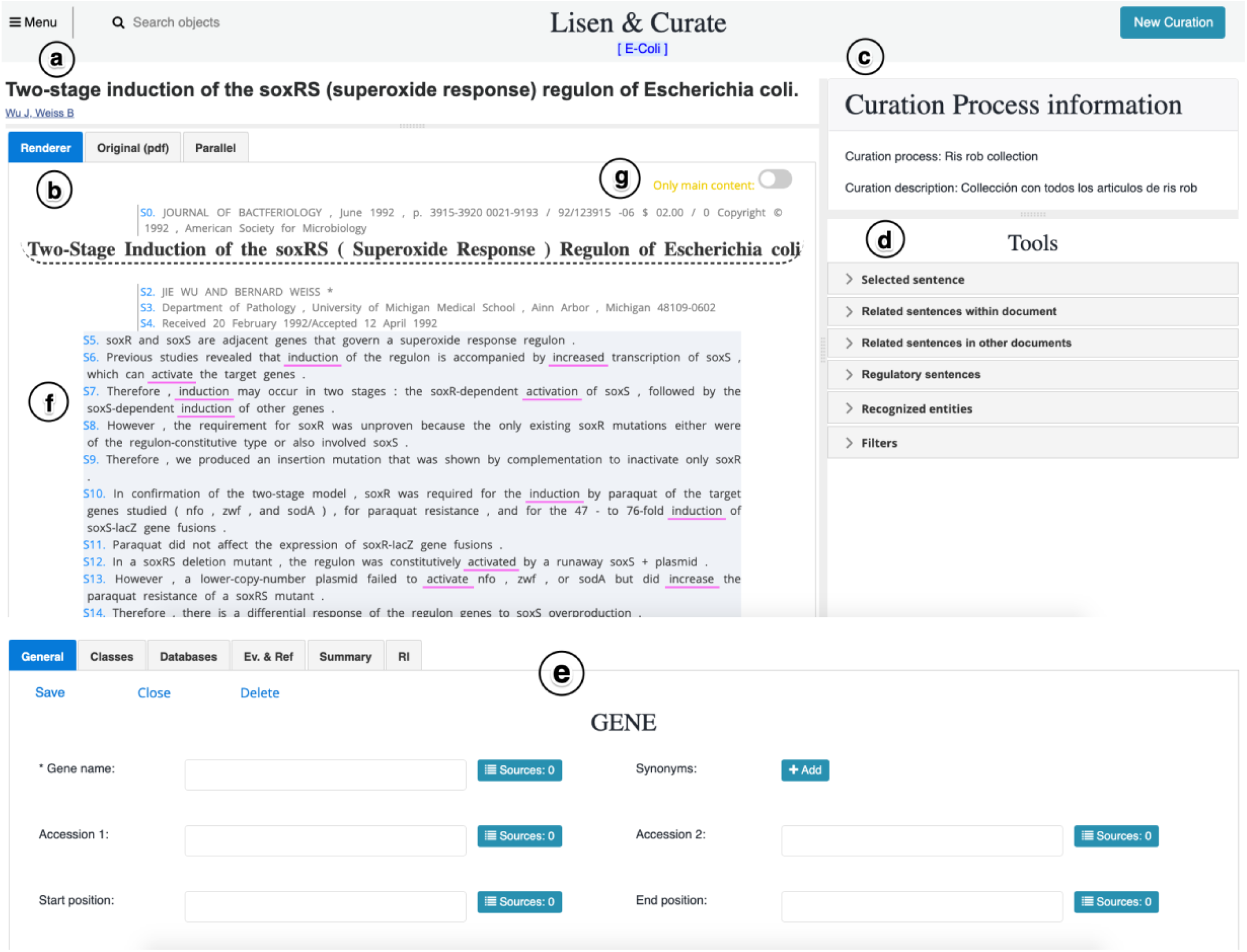
Main interface of the working environment.

Curators work with a structured and friendly visualization of articles, which is displayed using the JSON documents with sections and metadata obtained in the processing step. Articles could be visualized as text or PDF formats, with the option to display both formats side by side. Whether the article is shown as text format, this is presented as a list of numbered lines (Fig. 4f). Our tool to extract text from PDF files separates the main content of articles from secondary content, such as journal name, author names, headings, footers, etcetera. Secondary content is hidden while button *Only main content* is enabled (Fig. 4g), giving a *cleaner* visualization to curators. Nevertheless, if curators need to see secondary content, this appears by disabling the button.

#### Toolboxes

The *Tools* section of the main interface features a set of toolbox interfaces that present automatically extracted and enriched information to assist curation work. This information is the output of several text mining tools that are described in the *Tools for assisted curation* section. These toolboxes work only when the article is rendered as text format.

### Tools for assisted curation

#### Highlight and filter biological entities

For a while, curators of the RegulonDB team have employed the text mining tool OntoGene/ODIN (Rinaldi, 2013) to assist curation of regulatory interactions. The idea has been to select sentences that may contain regulatory interactions between genes and transcription factors (TFs) by filtering those sentences that contain a specific combination of tagged biological named entities and regulatory verbs (effects). This strategy was successfully applied to curate regulatory interactions for OxyR in E. coli K-12 (Gama-Castro, 2014).

We replicate a similar functionality for Lisen&Curate. JSON documents with recognized entities produced by processing step are employed to highlight entities and filter sentences that contain them. These entities are underlined with a defined color in sentences.

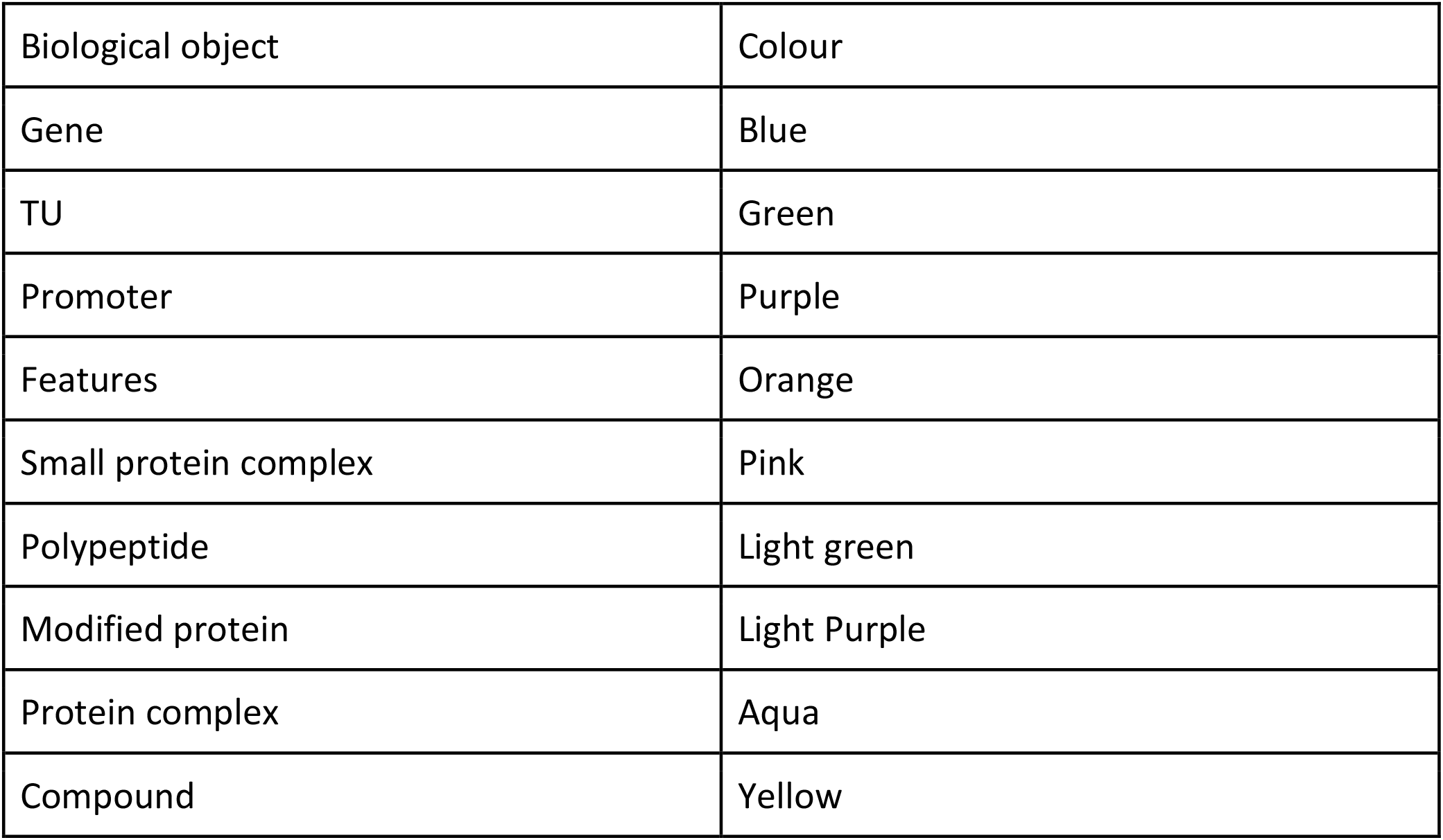
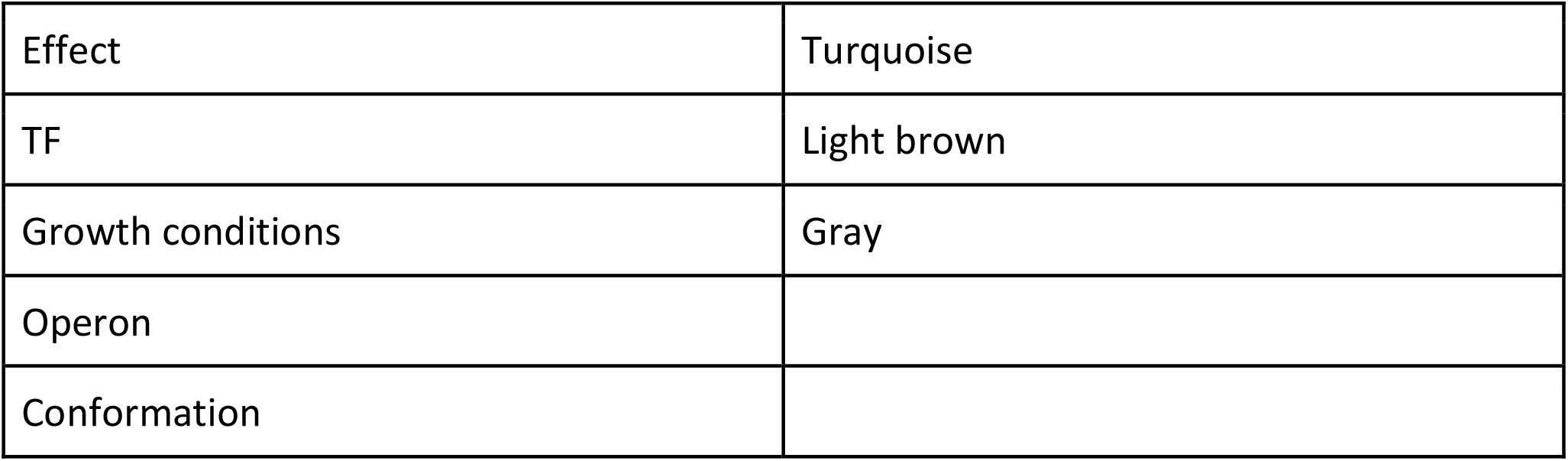

| Biological object | Colour |
| --- | --- |
| Gene | Blue |
| TU | Green |
| Promoter | Purple |
| Features | Orange |
| Small protein complex | Pink |
| Polypeptide | Light green |
| Modified protein | Light Purple |
| Protein complex | Aqua |
| Compound | Yellow |
| Effect | Turquoise |
| TF | Light brown |
| Growth conditions | Gray |
| Operon |  |
| Conformation |  |

In addition, entities and entity types are set in a table displayed by the *Recognized entities* tool. This table is useful to keep track of all the entities presented in the actual article. If curators select some entities in this table, the associated words in sentences are highlighted (Fig. 5).

**Figure 5.**
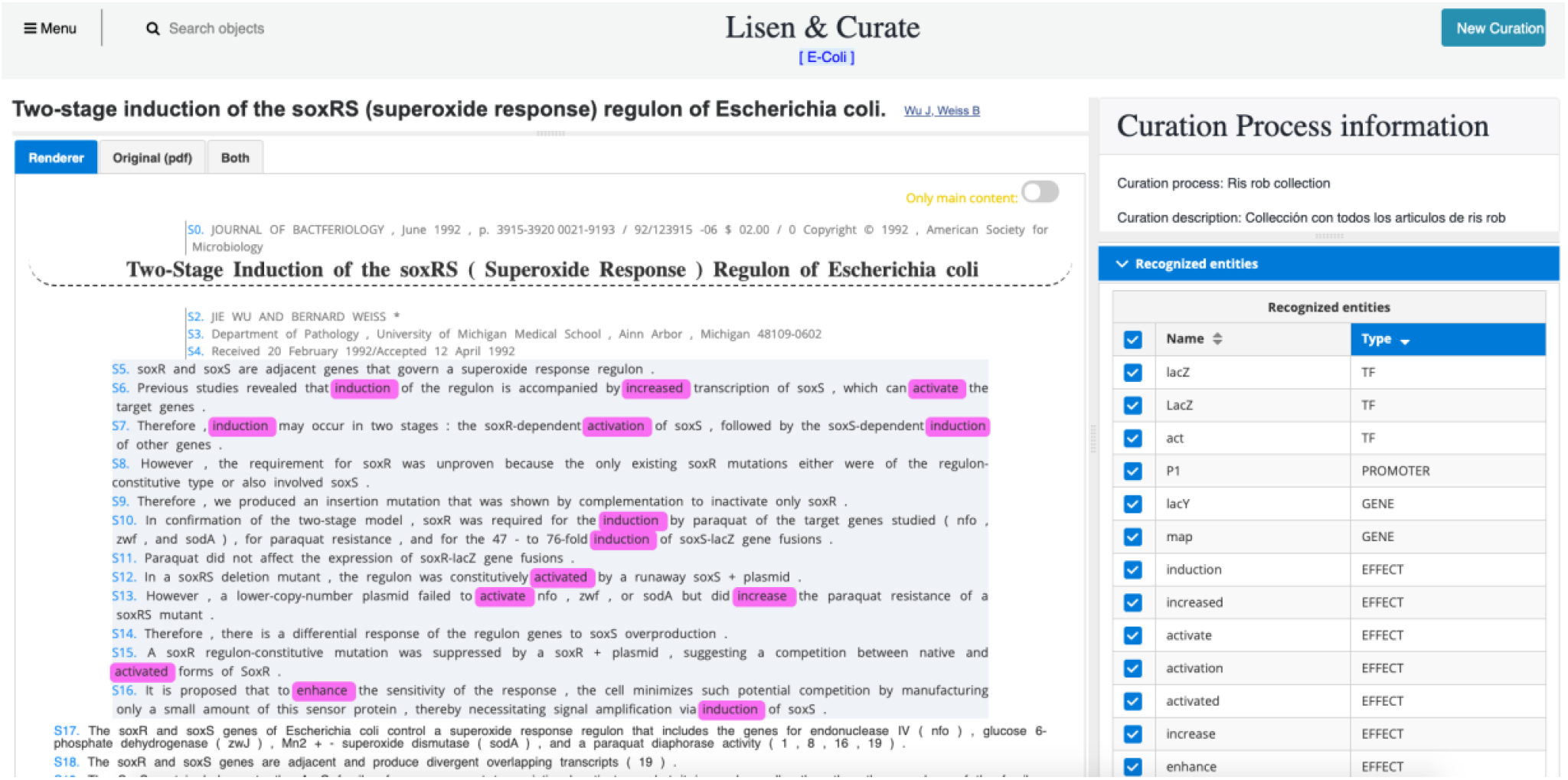
Recognized entities tool. Regulatory verbs (effects) selected in the table at right are highlighted in sentences at left.

#### Highlight regulatory sentences

One aim of Lisen&Curate is to display enriched information for curators to find relevant regulatory knowledge. Then, we included in Lisen&Curate a tool to highlight sentences with regulatory information (regulatory sentences). Using a curated data set of sentences containing regulatory interactions produced by the RegulonDB team (Santos-Zavaleta, 2019), we trained a predictive model to detect regulatory sentences. With this tool, we want to boost the speed of identifying undiscovered RIs within articles.

There are two advantages over detecting regulatory sentences using filters based on recognized entities. First, a predictive model may detect a regulatory sentence that contains a non-recognized entity, since this kind of models establish a decision region to separate sentences in two regions (classes) on a vector space: sentences with regulatory information from sentences without regulatory information. Then, a new sentence is classified according to the region it occurs, in spite of the entities it contains. Second, a predictive model offers the probability that a sentence pertains to a class, then, we can rank sentences by relevance using this probability.

The toolbox of Regulatory sentences shows a list with two fields: the number of regulatory sentences and the probability (Fig. 6). Curators can select a sentence from the list, and this will be highlighted in the article. We utilize a color gradient associated to the range of the probability, then higher probabilities are more colorful than the others. Highlighting regulatory sentences may allow curators at a glance to recognize the *level* of regulatory information that an article contains, so they may prioritize articles for curation.

**Figure 6.**
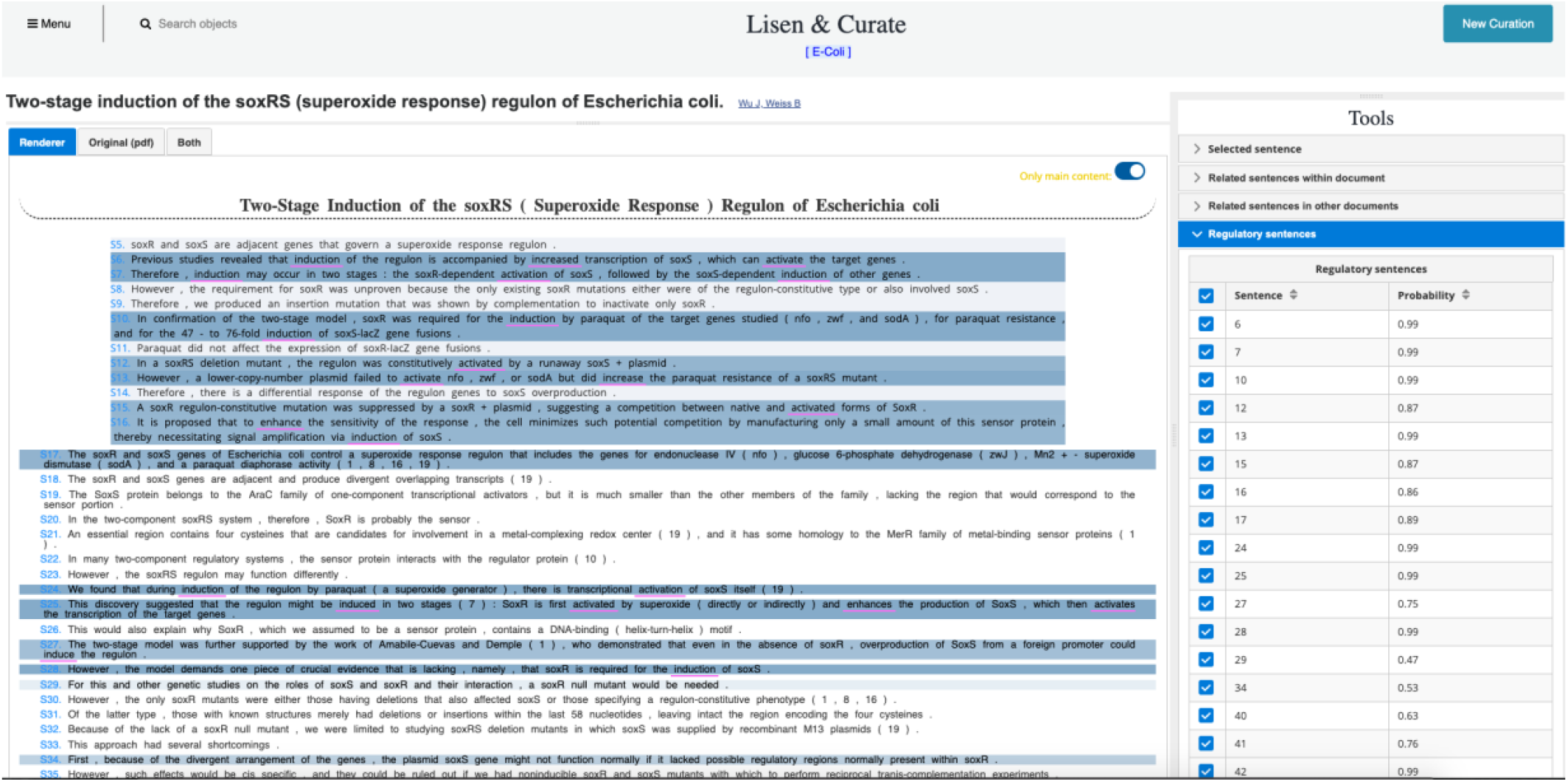
Highlight regulatory sentences. The color gradient is associated with the probability that the sentence is a regulatory sentence.

#### Curate extracted regulatory interactions

As we stated, a relevant motivation to develop Lisen&Curate was to expand the curation work of the RegulonDB team to literature of new bacteria. To face this challenge, we coupled to Lisen&Curate a text mining tool to extract RIs from literature. This is an in-house developed tool that extracts the regulator (TF), the regulated (gene or transcription unit), and the regulatory effect (regulation, activation, repression).

Our tool utilizes the linguistic concept of thematic roles. Thematic roles refer to the various roles that a nominal sentence plays with respect to the action described by a verb. Then, three constructions are used to extract RIs: active voice (ArgP activates argO, FimZ regulates its own transcription), passive voice (argO is activated by ArgP), deverbals (activation of rhaBAD by RhaS, Rob-regulated target inaA).

To extract RIs, text format files obtained in the processing step of articles are processed by our tool. The output is a JSON file with all extracted RIs. This file is incorporated to Lisen&Curate by a process that validates if the names of genes/TUs and transcription factors match with BioCyc identifiers. Then, extracted RIs are available for curator to review in a list (Figure 7). Users can display all RIs or only those of a curation process.

**Figure 7.**
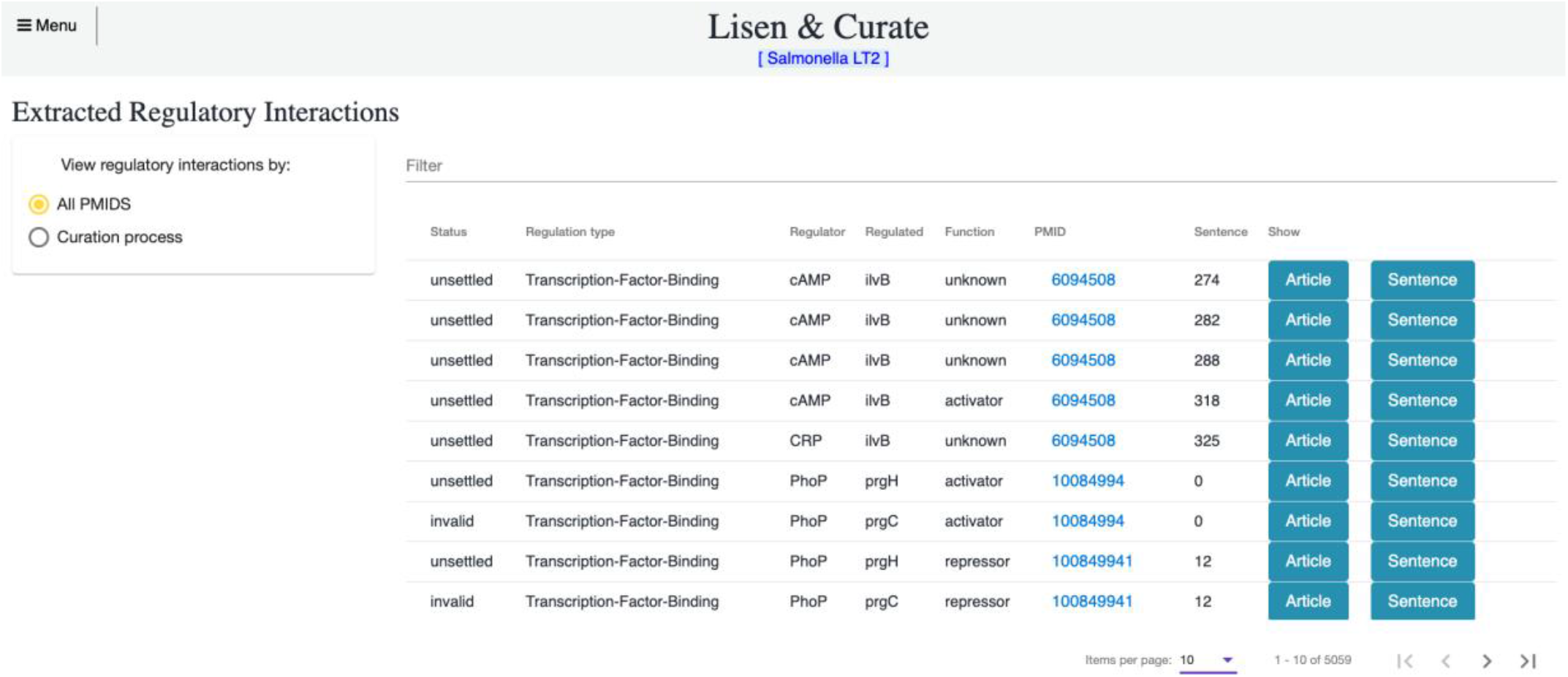
List of extracted RIs incorporated to the system.

In this list, information about the RI is showed: Status, Regulation type, Regulator, Regulated, Function (activator, repressor, unknown), Pubmed ID (PMID) and the number of the sentence where the RI is found. For the curation workflow in Lisen&Curate, RIs may have one of the following status:

- Unsettled: The RI is incorporated to Lisen&Curate and the names of the entities involved in the extracted RI match with BioCyc identifiers. Curators may change the status to Accepted, Modified, Rejected or Doubt.
- Invalid: The RI is incorporated to Lisen&Curate, but any of the names of the entities involved in the extracted RI do not match with BioCyc identifiers. Curators have to manually correct identifiers.
- Accepted: All the data involved in the extracted RI is correct and the related sentence is a reliable textual source of evidence.
- Modified: Data of the extracted RI were corrected by curators and the related sentence is a reliable textual source of evidence.
- Rejected: Sentence related to extracted RI does not give reliable textual source of evidence of the RI.
- Doubt: Sentence related to extracted RI does not give enough information for curator to decide if the RI is incorrect or incorrect. The RI cannot be accepted or rejected until more information is provided.

Curator can access directly to the article that contains the extracted RI using this list (button *Article*). Then, the article is displayed in the main interface and, automatically, the system points to the sentences within the article. Also, the extracted RI is selected in the toolbox of extracted RIs of the article (Figure 8).

**Figure 8.**
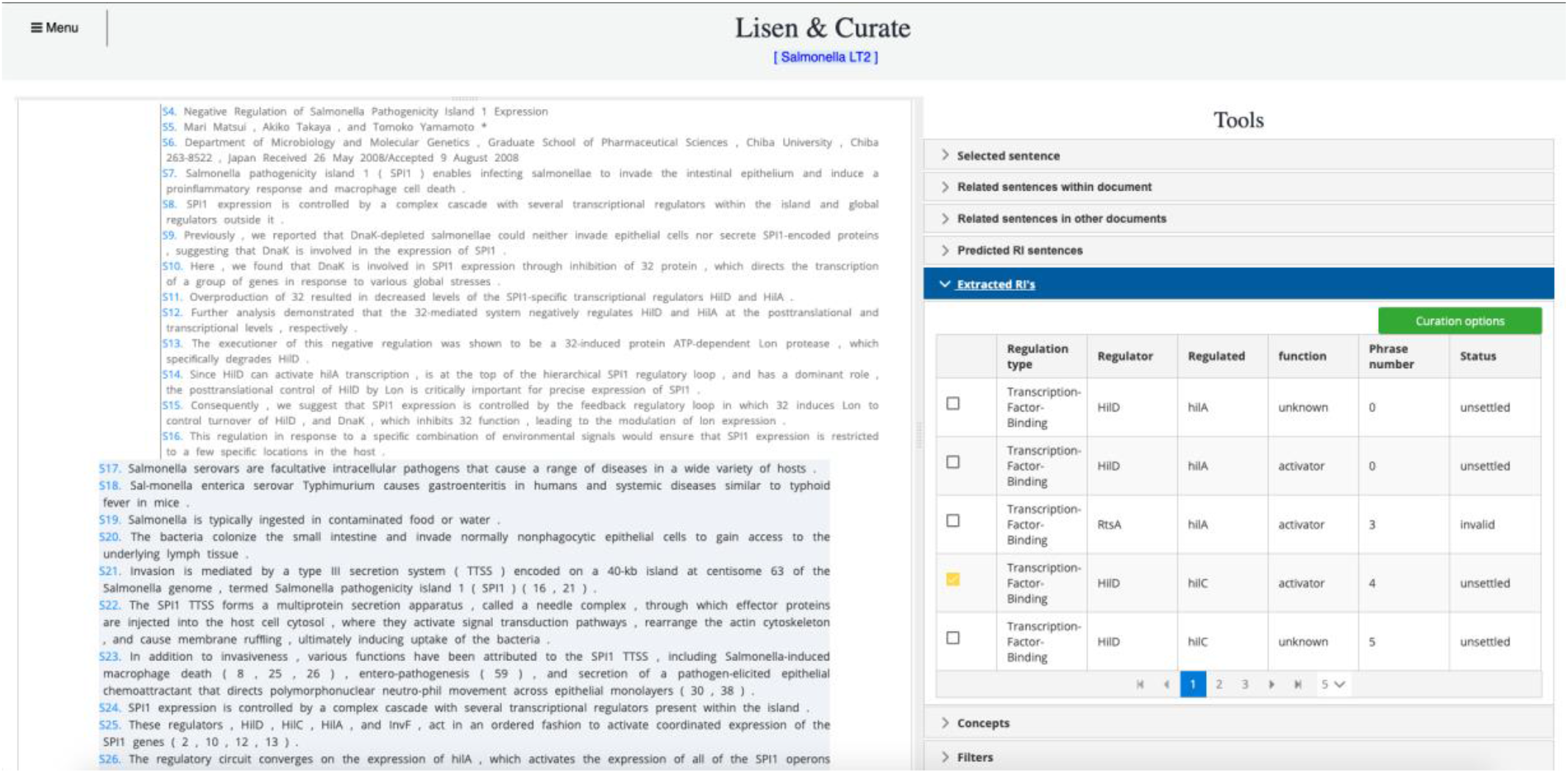
Main interface with toolbox of Extracted RIs. At the right, the toolbox shows the list of extracted RIs from the article. The curator can modify the status of a RI using the button *Curation options*.

Using this toolbox, curators can change the status of an RI after checking the textual source of evidence (Figure 9). The status Accepted and Modified will store the data of the extracted RI in our database. Automatically, the system also stores the sentence associated to the RI as a textual source of evidence that will be published in our database systems (RegulonDB, L-Regulon). Also, RIs classified in Accepted, Rejected and Doubt will be used to improve text mining systems to extract RIs, since they are negative and positive examples. From a machine learning point of view, they are training data.

**Figure 9.**
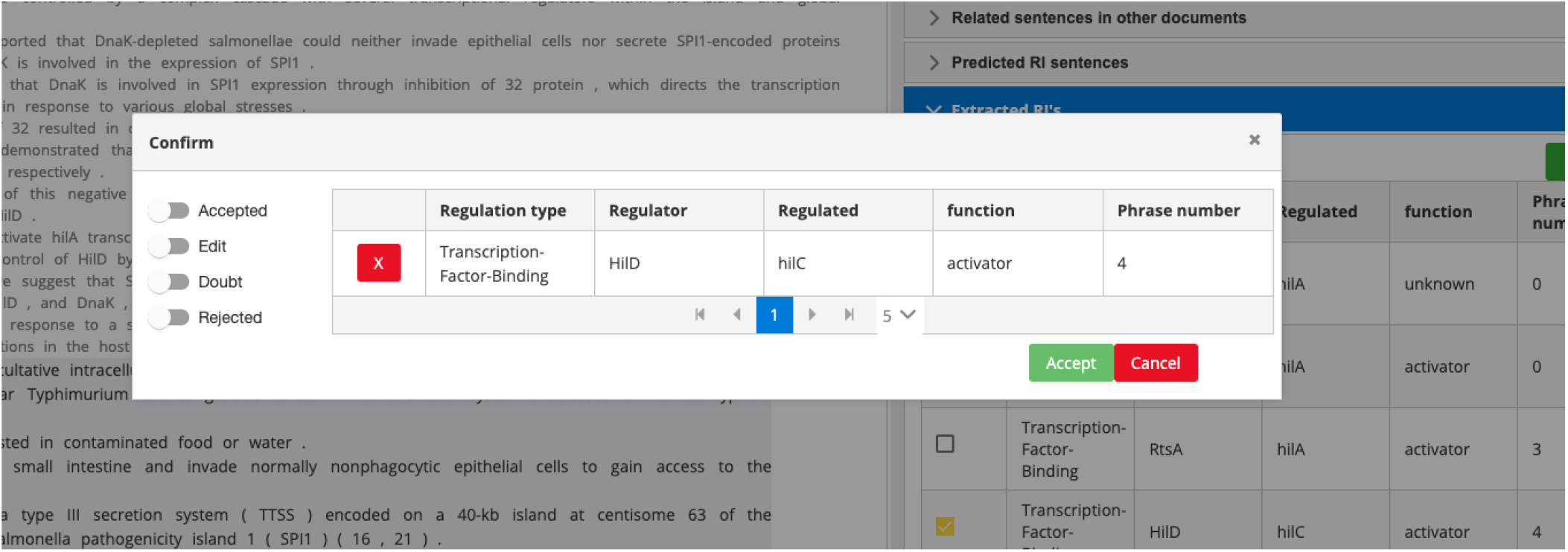
Interface to change the status of extracted RIs.

#### Interfaces for capturing data

Interfaces for capturing data aim to facilitate the register of regulatory data of different biological objects. Captured data are stored in the local database of the organism. Lisen&Curate comprises interfaces to capture and query data for the following objects:

- Gene
- Transcription units
- Promoters
- Regulatory interactions
- Proteins
  - Polypeptides
  - Modified protein
  - Small molecule complex
  - Protein complex
- Protein compounds
- Protein features
- Terminators

All interfaces for capturing data have the same appearance. They are divided into tabs for better organization. Each tab contains the necessary fields for recording regulatory data. As an example, we show in Figure 10 the interface for capturing data of genes. An innovative feature of Lisen&Curate is that each field can be associated to the textual source where the field is described (button *Sources*). This textual source may be the complete sentence or just a fragment.

**Figure 10.**
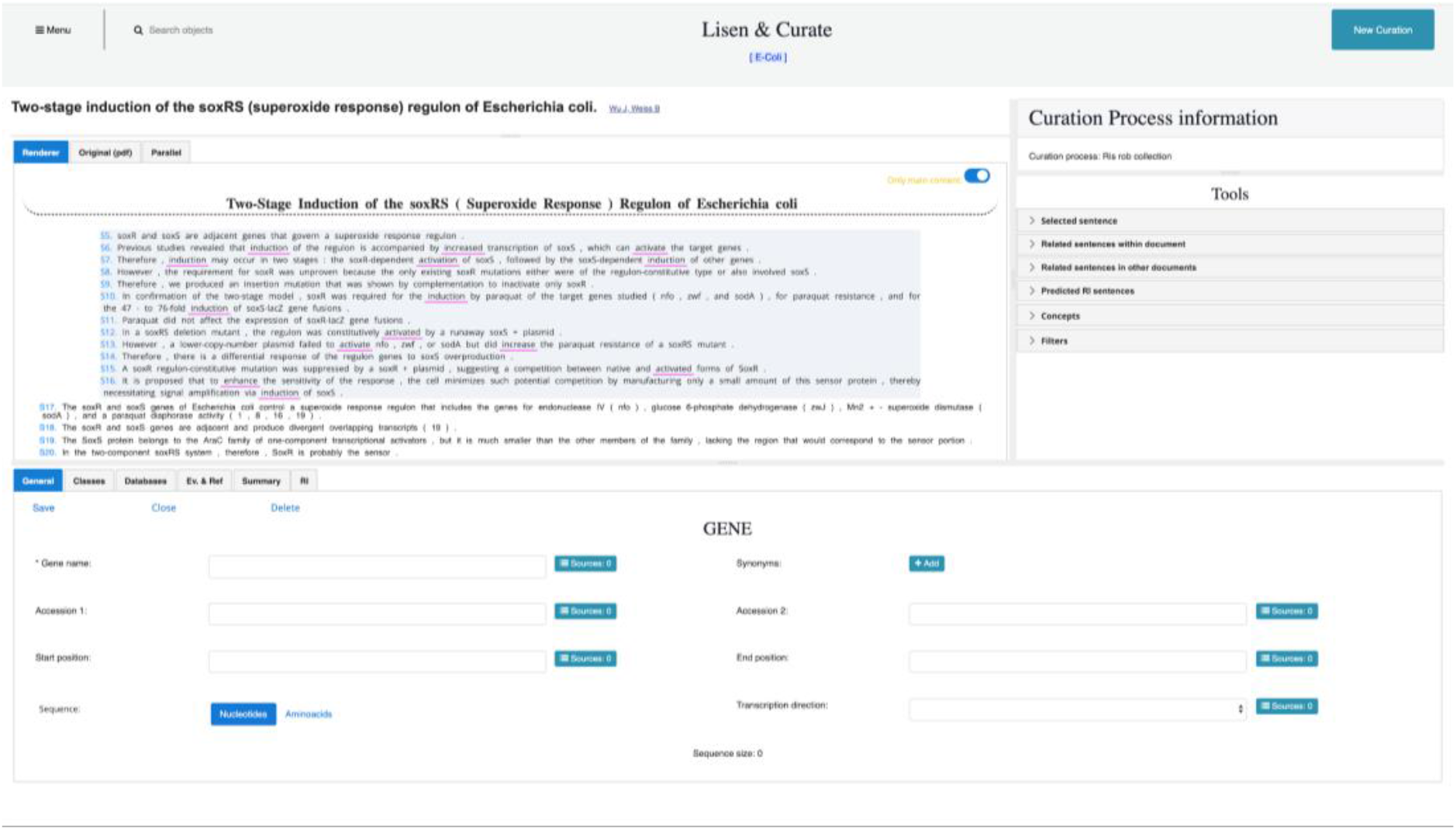
Interface for capturing data of genes.

All interfaces share the main three tabs:

- General: To add and display the core information of the object.
- Ev. & Ref.: To add and show sources of evidence to support the stored regulatory data of the object.
- Summary: To add and display additional information that is not covered by the fields of the General tab.

Depending on the object, there are also the following tabs:

- Databases: To add and show the links of the object to external databases.
- Classes: To add and display the class of the object from the BioCyc ontology.
- RI: To add and show the regulatory interactions where the object participates.
- TU: Appears in the Promoter interface to add and show the transcription units where the promoter participates.
- Promoter: Appears in the Transcription unit interface to add and show a promoter.
- Promoter boxes: Appears in the Promoter interface to add and show promoter boxes.
- Terminator: Appears in the Transcription unit interface to add and show a terminator.
- Reactions: Appears in Compound interface to add and show the produced reactions.
- Genes: Appears in the interfaces of the four types of proteins to add and show genes.
- GO: Appears in the interfaces of the four types of proteins to add and show growth conditions.
- Features: Appears in the Polypeptide interface to add and show the protein features.
- Other proteins: Appears at three of the four protein interfaces (Polypeptide is excluded) to add and show the other forms of the protein.
- Subunits: Appears in the Protein complex interface to add and show subunits.
- Components: Appears in the Protein small molecule complex interface to add and show components.

Lisen&Curate consumes data from BioCyc to display the information of objects. BioCyc is an available resource including 14,735 Pathway/Genome Databases (PGDBs) for describing several organisms and tools for obtaining related data (https://biocyc.org/) (Karp, 2019). Users of Lisen&Curate can search BioCyc information of objects using a search bar located at the top of the main interface.

#### Reporting curation work

In Lisen&Curate, operations to add or modify database records are logged to generate reports of curation workload. Two reports are available:

- A bar plot to visualize the number of biological objects added or modified in the system within a defined period.
- A downloadable report generated as a spreadsheet with the detailed information of a list of specific objects within a defined period.

## METHODS

Lisen&Curate was developed with a robust and flexible software architecture and several technological components to deal with structured and unstructured information (Figure 12). To consume data from BioCyc we employed Pathway-Tools (Karp, and others, 2016) using PythonCyc, which provides the basic functions for accessing Pathway/Genome Databases (PGDBs). The PythonCyc package is hosted on GitHub (https://github.com/ecocyc/PythonCyc) and is a separated installation from Pathway-Tools.

**Figure 12.**
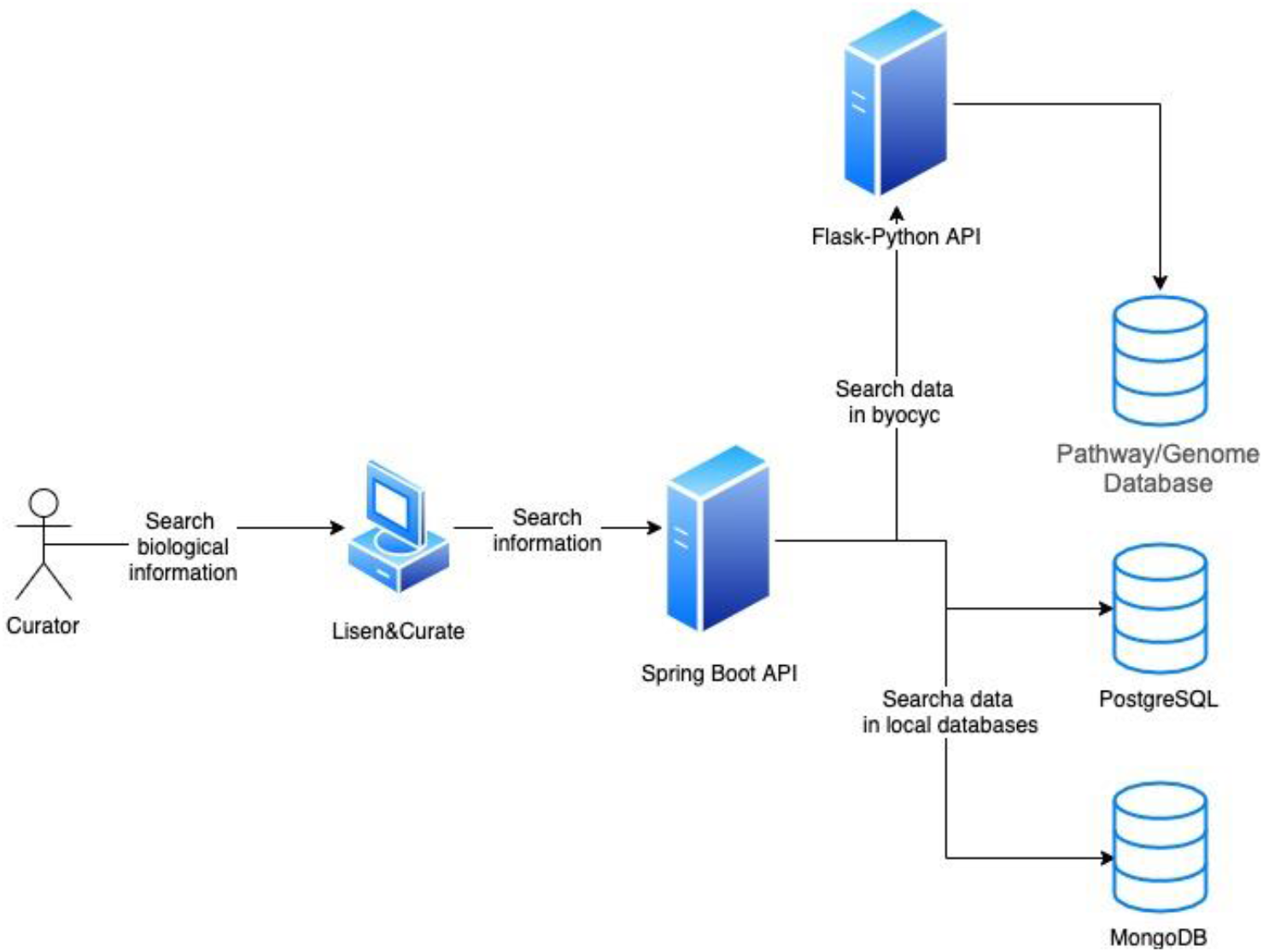
Lisen&Curate software architecture.

We coded with Python version 2.7 the REST type web services to extract information from Pathway-Tools and we use Flask, a lightweight WSGI web application framework (https://palletsprojects.com). We employed Spring Boot (https://spring.io/projects/spring-boot) as an infrastructure for the development of Spring-based Full Rest Web services.

As Lisen&Curate process structured and unstructured data, we used different databases technologies. To store textual data we utilized MongoDB (https://www.mongodb.com), which stores data in flexible, JSON-like documents, with meaning fields that can vary from document to document whose data structure can be changed over time. Structured data related to regulatory information is stored in PostgreSQL (https://www.postgresql.org). This is a powerful open source relational database that extends the SQL language and gives a secure and scalable store of demanding data workloads. Finally, we used Elasticsearch (https://www.elastic.co), an open source distributed analytic and analysis engine for search tagged entities within JSON files produced by processing step.

## ACKNOWLEDGMENTS

We acknowledge technical support by Luis José Muñiz-Rascado, Víctor Del Moral and César Bonavides-Martínez, as well as Cecilia Ishida-Gutiérrez and Víctor Tierrafría for their input in using the system. J.C.-V. acknowledges support by DGAPA from UNAM during his sabbatical leave at the Center for Genomic Regulation in Barcelona.

